# Rif-Correct, a software tool for bacterial mRNA half-life estimation

**DOI:** 10.1101/2022.02.25.481994

**Authors:** James R. Aretakis, Jared M. Schrader

## Abstract

**Background:** Genome-wide measurement of bacterial mRNA lifetimes using the antibiotic rifampicin has provided new insights into the control of bacterial mRNA decay. However, for long polycistronic mRNAs, the estimation of mRNA half-life can be confounded with transcriptional runoff caused by rifampicin’s inhibition of initiating RNA polymerases, and not elongating RNA polymerases.

**Results:** We present the Rif-correct software package, a free open-source tool that uses transcriptome models of transcript architecture to allow for more accurate mRNA half-life estimates that account for the transcriptional runoff. Rif-correct is implemented as a customizable python script that allows for users to control all the analysis parameters to achieve improved mRNA half-life estimates.

**Conclusions:** Rif-correct is the first free open-source computational analysis pipeline for Rif-seq dataset mRNA half-life estimation. It is simple to run, fast, and easy to run with a detailed instruction manual and example datasets.

## Background

mRNA half-lives can be measured genome-wide in bacteria using the antibiotic Rifampicin to shut of mRNA transcription and measuring abundance at different time-points by microarray or RNA-seq (1,2) (Fig. 1 A., Rif-Seq description). While generally successful, complexities arise from the antibiotic rifampicin, which only blocks transcription initiation, and does not inhibit transcription elongation (3) (Fig 1b). As a result of the rifampicin mode of action, some elongating RNA polymerases (RNAP) can continue to transcribe and run off. Therefore, directly fitting a simple half-life equation to the natural log transformed mRNA abundances does not accurately fit the data, especially for long operons, causing an inaccurate determination of apparent half-life from the slope (Fig 1 C.). To overcome this expensive proprietary software has been used to correct their data for transcriptional runoff (1). Therefore Rif-Correct was developed as free open-source software that uses Rif-seq data and RNAP elongation rate to improve the mRNA half-life estimates by correcting for the mechanism of action issues inherent to rifampicin.

**Figure 1:**
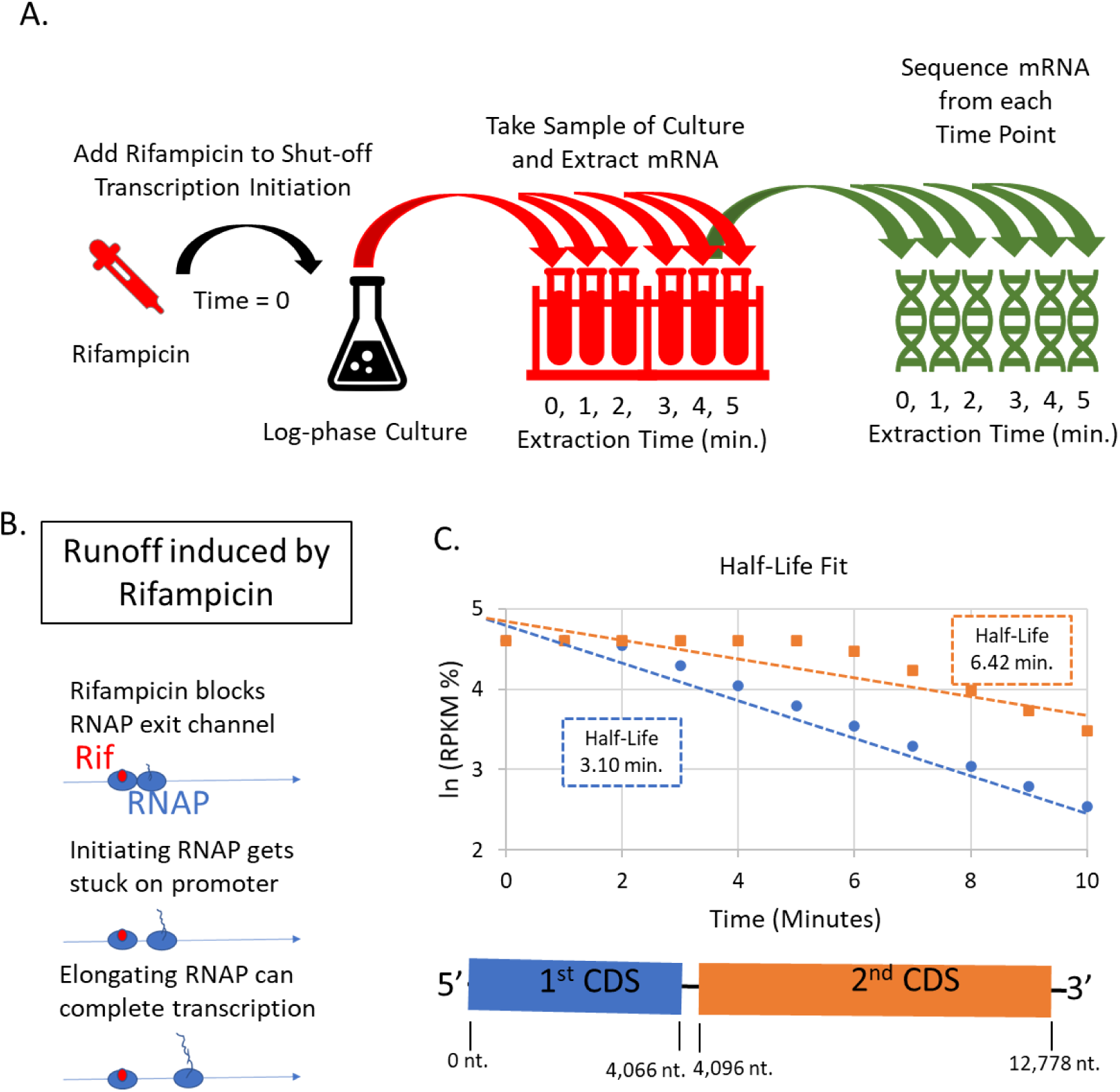
Using Rif-Seq to determine mRNA half-lives: A) The Methodology for the Rif-Seq experiment is shown. The antibiotic Rifampicin is added to a steady-state log phase bacterial culture at time = 0 to shut-off transcription initiation. Then samples of the culture are taken at different time points after the addition of Rifampicin and the cellular RNAs are quenched in an RNA protection solution. Finally, RNA-Seq libraries are made from each time point which report the relative genome-wide mRNA abundance (RPKM). B) Natural log transformed mRNA abundance (RPKM) for a two gene example operon with the blue circles representing the data for the first gene in the operon and the orange squares representing the data for the second gene in the operon. Linear regressions (shown as the blue and orange dotted line, respectively) does not fit the data well due to the two-phase decay resulting from runoff followed by decay. Importantly, both genes are part of the same mRNA with half-lives of 2.77 minutes. C) The antibiotic rifampicin’s mechanism of action inhibits RNAP transcription initiation by blocking the exit tunnel(3), but not transcription elongation where the exit channel is already occupied by RNA. Consequently, there is a runoff of elongating RNAP that can cause a delay to the observed decay of the mRNA which is the source of poor curve fitting.

## Implementation

### Inputs

To run Rif-correct (available at https://github.com/schraderlab/rif_correct) the user needs to input multiple data files (Fig 2 A). This includes 1) a tab delimited mRNA abundance file with mRNA abundance measurements at each time point after rifampicin collection together with the name for each mRNA. For genome-wide analysis, RNA abundances will be measured by RNA-seq or by microarray. 2) a tab delimited genome annotation file containing the locus tag for each coding sequence with genomic coordinates and strand information, the operon number and list of genes within each operon. This genome annotation file should be ordered with positive stand in genome order from origin to terminus, and then negative strand from terminus to origin. To arrange the genome annotation file in this manner, a small standalone python script “CDS order” is provided to perform this step. 3) an optional transcriptome file can be added to take full advantage of the runoff correction for operons it is needed. This file is an assembly of the CDS, transcription start site (TSS) data, and operon structure to build probably transcriptional units. This file includes: If CDS with that operon number are in a polycistronic operon, start gene of operon, last gene of operon, strand, farthest upstream promoter site (necessary) and up to 3 other promoter sites (optional), TSS type (optional), number of TSSs (optional), internal TSS (optional), a flag as to whether an operon is simple (single mRNA isoform predicted) or complex ((optional, multiple mRNA isoforms predicted), number and names of internal non-coding RNAs to operon (optional). A complete set of instructions to generate each file and example files are included.

**Figure 2:**
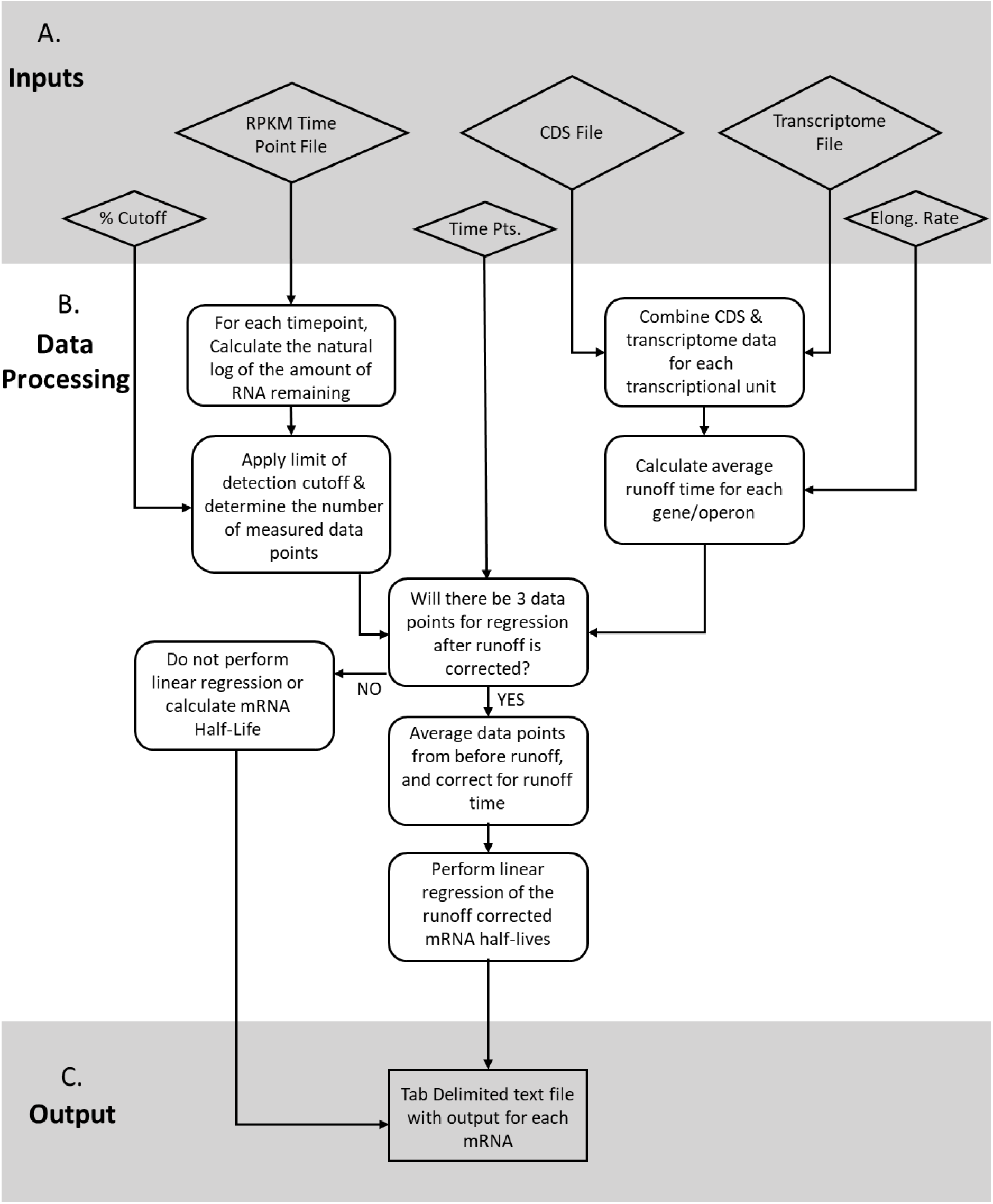
Rif-Correct software workflow A) Inputs for the Rif-Correct Software is shown. Large diamonds denote input files for Rif-correct, smaller diamonds denote input values typed into the command line by the user. B) Data processing steps and decisions for the Rif-Correct software is shown. C) Output for the Rif-Correct Software is a tab delimited text file. The output file contains information from each inputted mRNA.

In addition to these data files, the user is also prompted to enter multiple inputs for the data at the command prompt (Fig 2 A). First, the user must input the time points after rifampicin addition in which the RNA abundance measurements were collected. Second, the user must input the elongation rate for RNAP on mRNA in nucleotides per second (An example for *E. coli* ranges from 41nt/sec in M9G to 55 nt/sec in LB (4), and *C. crescentus* is 19 nt/sec in M2G media (Schrader lab unpublished data)). The lower limit of detection to be used in mRNA abundance estimation (optional), expressed in the percent of the original mRNA level remaining (3% is recommended). The lowest acceptable correlation value from linear curve fitting that to be acceptable for reporting of a half-life (R > 0.7 recommended, optional).

### Data Processing

The Rif-correct workflow is shown in Figure 2B. First, the genome annotation file and the optional transcriptome file are combined into one data structure if provided. This creates one master list of all predicted transcriptional units with their corresponding information, but if the transcriptome file is not provided the list is populated with null values and the program will treat each gene as monocistronic and the start codon will be used instead of the TSS for transcription runoff time estimation. The transcription runoff time is calculated for each transcriptional unit based on the average elongation rate of the RNAP on mRNA, if the transcriptome file is not provided, or the elongation rate is set to 0 nt/sec then there will be no runoff correction for any gene. To calculate runoff time, RNAP elongation time is calculated to the middle of each CDS feature. This distance is calculated from half the distance from the farthest upstream TSS site, or first CDS start codon if no TSS is known (for operons this would be the first gene in the operon), to the end of the CDS site for that specific gene. For polycistronic operons, this runoff time is estimated for each CDS on each transcriptional unit. The mRNA abundance measurement data for each CDS and time point is calculated as a percent based on its level compared to the 0-minute time point, and then the natural log of this ratio is calculated. If the user has placed a limit of detection cutoff, any data points below the cutoff are discarded from further analysis. To account for the transcription runoff time, for each CDS any RNA abundance data points before the runoff time are averaged together and included as a calculated data point with the estimated runoff time. The number of consecutive time points with RNA abundance measurements are then recorded for each CDS. Next, linear curve fitting is performed to calculate the Rif-corrected mRNA half-life. For curve fitting of mRNA half-lives, there must be at least three consecutive data points after runoff. If there are not enough data points for a CDS, then no linear regression is performed, and no half-life is calculated for that gene and the data for the gene is written to the output file. If there are enough data points for a CDS then no calculation of mRNA half-life will be calculated. If the correlation value of curve fitting is below the optional curve fit threshold for a given CDS, no mRNA half-life will be calculated. Finally, a tab delimited text file is saved (Fig 2 C).

## Output

A tab delimited text file with the user inputted name is created in the directory where the program is run. This file contains: Slope of the Linear regression of the natural log transformed and transcription run off corrected data, Y intercept of the Linear regression of the natural log transformed and transcription run off corrected data, R value of the Linear regression of the natural log transformed and transcription run off corrected data, Rif-corrected half-life in minutes based in the slope (ln(2)/-slope), number of datapoints used in linear regression after the low abundance threshold cutoff was applied, Number of data points averaged due to the runoff correction, R value flag if the linear regression R value is above the user inputted R value cutoff (R = -1 is perfect natural log decay, R = 0 is a flat line, and R = 1 is an perfect increasing half-life), Too slow to measure mRNA half-Life flag - if the calculated half-life is negative or twice as long as the last time point used in the linear regression, and there are also quality control outputs reported for each mRNA.

## Results/Discussion

Rif-Correct allows the use of transcriptome data for models of mRNA architecture (or from the CDS +1 site if the TSS is not known) to be used to correct for transcription runoff induced by rifampicin. While many short mRNAs do not require any correction for half-life estimation, long polycistronic operons are aided significantly by the Rif-correct algorithm (Fig 3). For example, hypothetical mRNA abundance data were generated for a hypothetical simple 2-gene operon that is degraded with a 2.77 minute half-life, and half-lives of each CDS region were fit to both a simple half-life equation (Fig 1C) or by the Rif-correct package (Fig 3). Despite being part of the same mRNA with the same half-life, simple half-life fitting yields a half-life of 3.10 min for the first CDS, and 6.42 min for the second CDS, while Rif-Correct runoff correction yields 2.77 min for the first CDS and 2.77 min for the second CDS. Overall, this example highlights the runoff artifact where longer mRNAs appear to have artificially extended mRNA lifetimes by simple half-life fitting. When Rif-correct is applied transcriptome wide, such analysis can help to ensure more accurate mRNA half-lives are obtained.

**Figure 3:**
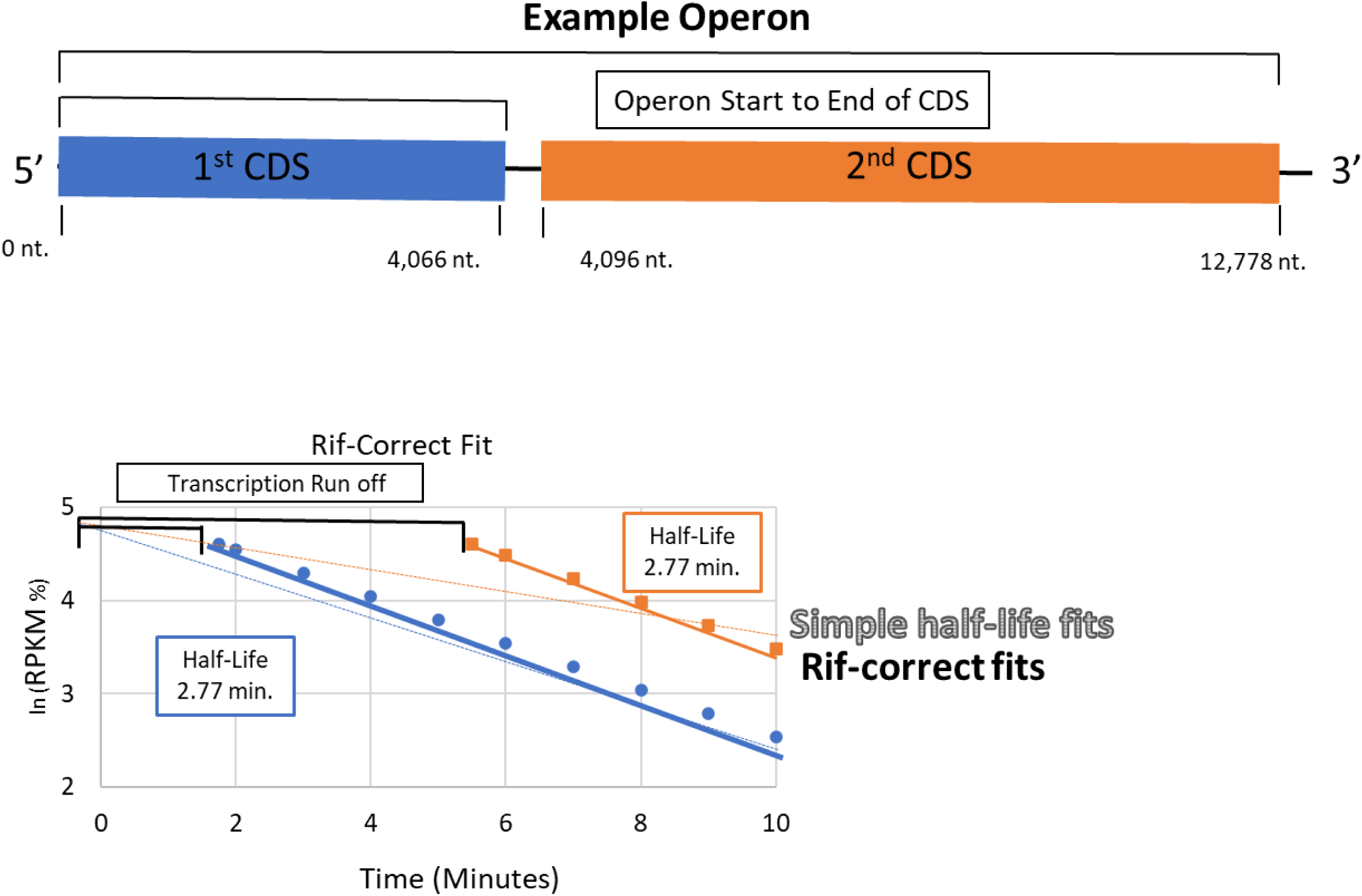
Rif-Correct Software Fit The example operon data from figure 1 is shown with the 1^st^ CDS region in blue, and the 2^nd^ CDS region in orange. Bracketed lines represent the operon start to end of CDS used in the transcription runoff correction. Rif-Correct software improves curve fitting and corrected for the transcription runoff. Natural log transformed relative RPKM levels are shown for the two gene example operon with the blue circles representing the data for the first gene in the operon and the orange squares representing the data for the second gene in the operon. Linear regressions are shown for the simple half-life fits (shown as the blue and orange dotted line), and for the Rif-Correct fit (shown as the blue and orange solid line). The difference in slope between the Rif-correct fit (2.77 min, and 2.77 min) and the simple fit (3.10 min, 6.42 min) show how using the Rif-Correct Fit, allows for a more accurate half-life calculation (half-lives shown in the blue and orange solid boxes, respectively).

## Conclusions

Rif-Correct is a free open-source software that uses a transcriptome-based model to correct for transcriptional runoff to improve mRNA half-life estimates in bacteria. It is highly customizable with parameters to allow researchers to perform the analysis in the best way for their applications and is fast and easy to run. With a growing number of Rif-seq datasets, Rif-correct will be broadly useful to bacterial mRNA decay researchers to help them to calculate mRNA half-lives more easily across the transcriptome.

